# The Cellular and Extra-Cellular Proteomic Signature of Human Dopaminergic Neurons Carrying the LRRK2 G2019S Mutation

**DOI:** 10.1101/2024.09.20.614055

**Authors:** Felix Knab, Giambattista Guaitoli, Mohamed Ali Jarboui, Felix von Zweydorf, Fatma Busra Isik, Franziska Klose, Anto Praveen Rajkumar, Thomas Gasser, Christian Johannes Gloeckner

## Abstract

Extracellular vesicles are easily accessible in various biofluids and allow the assessment of disease-related changes of the proteome. This has made them a promising target for biomarker studies, especially in the field of neurodegeneration where access to diseased tissue is very limited. Genetic variants in the LRRK2 gene have been linked to both familial and sporadic forms of Parkinson’s disease. With LRRK2 inhibitors entering clinical trials, there is an unmet need for biomarkers that reflect LRRK2-specific pathology and target engagement. In this study, we used induced pluripotent stem cells derived from a patient with Parkinson’s disease carrying the LRRK2 G2019S mutation and an isogenic gene corrected control to generate human dopaminergic neurons. We isolated extracellular vesicles and neuronal cell lysates and characterized their proteomic signature using data-independent acquisition proteomics. We performed differential expression analysis and identified 595 significantly differentially regulated proteins in extracellular vesicles and 3205 in cell lysates. Next, we performed gene ontology enrichment analyses on the dysregulated proteins and found close association to biological processes relevant in neurodegeneration and Parkinson’s disease. Finally, we focused on proteins that were dysregulated in both the extracellular and cellular proteomes and provide a list of ten promising biomarker candidates that are functionally relevant in neurodegeneration and linked to LRRK2 associated pathology. Among those was the sonic hedgehog signaling molecule, a protein that has tightly been linked to LRRK2-related disruption of cilia function. In conclusion, we characterized the cellular and extracellular proteome of dopaminergic neurons carrying the LRRK2 G2019S mutation and propose an experimentally based list of promising biomarker candidates for future studies.

## INTRODUCTION

Parkinson’s disease (PD) is a neurodegenerative disorder affecting millions of patients worldwide and causing both significant morbidity and impaired quality of life (*1*). A combination of genetic and environmental factors has been implicated to trigger the PD pathogenesis (*2*). Among the genetic factors associated with PD, variants in the leucine-rich repeat kinase 2 (LRRK2) gene have garnered significant attention. The *LRRK2* G2019S mutation represents the most common genetic cause of familial PD (fPD)(*3,4*) and can also be found in sporadic PD (sPD) patients (∼2%) due to reduced penetrance of the mutation (*3–5*). Genome-wide association studies (GWAS) have further revealed that frequent polymorphisms in the vicinity of LRRK2 are linked to an increased risk of developing sPD (Simón-Sánchez et al., 2009). Various potential roles for LRRK2 on the cellular level of the PD pathogenesis have been reported and are often associated with the increased kinase activity known to be caused by several of the described LRRK2 mutations (*6*).

Although PD is currently diagnosed clinically, the importance of protein based biomarkers is increasing (*7*). Analyses of protein markers such as Neurofilament Light Chain Protein or α-synuclein in the cerebrospinal fluid (CSF) or blood of patients are becoming more relevant and recent breakthroughs around α-synuclein seeding assays have the potential to re-shape the clinical routine within the next years (*8,9*). However, none of the biomarker assays approaching the clinical routine can pin down the intra-individual PD pathogenesis. With multiple potential LRRK2 inhibitors being developed and entering clinical trials, a biomarker that reflects LRRK2-specific molecular pathology, target engagement and cellular response is of great interest to the scientific community (*10*). With LRRK2 likely playing a role in some but not all sPD patients, identification of patient subgroups benefitting from inhibitor treatment would further represent an important starting point for the design of future clinical trials. Since its identification as a LRRK2 substrate, Rab proteins have been discussed and their phosphorylation has been utilized as proxies for LRRK2 kinase activity (*10,11*), but it can be assumed that mutated LRRK2 leads to more than just increased phosphorylation of direct target molecules. It is therefore important to broaden our portfolio of protein markers indicating LRRK2- related pathophysiology.

In the present study we aimed to provide an experimentally based list of LRRK2-related protein biomarker candidates. Such candidates should ideally be: (1) functionally relevant in the context of neurodegeneration, (2) functionally linked to LRRK2, (3) reflecting cellular pathogenesis, while also being (4) easily accessible. For this purpose, we used data-independent acquisition mass spectrometry (DIA-MS) to characterize the proteome of extracellular vesicles (EVs) isolated from induced pluripotent stem cells (iPSCs) derived human dopaminergic neurons (hDaNs) and direct cell extracts. This line had previously been generated from an iPSC-line derived from a PD patient carrying the LRRK2 G2019S mutation together with an isogenic gene corrected control (*12,13*).

EVs are membranous vesicles secreted by a variety of cell types into the extracellular space and next to nucleic acids contain intracellular proteins (*14*). They have proven to be easily accessible from biofluids such as plasma or CSF and throughout the last decade have been intensively studied as sources for biomarkers (*15,16*). We compared the EV proteome to the cellular proteome of the hDaNs to identify proteins that are dysregulated both on a cellular and extra-cellular level and to increase the robustness of our findings. Finally, we performed gene ontology analyses and identified LRRK2 interactors among the dysregulated proteins to pin down functionally relevant candidates.

## EXPERIMENTAL PROCEDURES

### Cultivation of Human Dopaminergic Neurons

We previously described the generation of patient-derived iPSCs and neuronal progenitor cells (NPCs) from two PD patients carrying the LRRK2 G2019S mutation (*12,13*). For this study, NPCs from one of the two patients, a female with an age of onset of 40, was used together with the respective gene corrected control line and will be referred to as L1 GC (gene corrected) and L1 G2019S, formerly referred to as L1. The induction of dopaminergic differentiation of NPCs together with the thorough characterization of the resulting human dopaminergic neurons (hDaNs) has recently been published (*17*)Briefly, for each genotype, one cryo-tube of NPCs was thawed and cells were split into three 6- wells. The three 6-wells were expanded independently and functioned as technical replicates for the isolation of EVs and cell lysates. NPCs were cultured and expanded in NPC media (Supplemental Table S1) until they reached 80% confluency, after which differentiation was induced using a media referred to as D7 media (Supplemental Table S1). After seven days, D7 media was exchanged for maturation media (Supplemental Table S1). On day 14 of the differentiation, 2 ml of maturation media was put into each 6-well of an entire 6-well plate. Conditioned cell culture media (CCM) was collected and pooled from one 6-well plate after three days, resulting in a total of 12 ml of CCM per technical replicate. This was repeated every three days until day 23 of the differentiation. CCM from each collection time point was pooled, resulting in a total of 36 ml per technical replicate and was processed further for isolation of EVs (Fig. 1).

**FIG 1.**
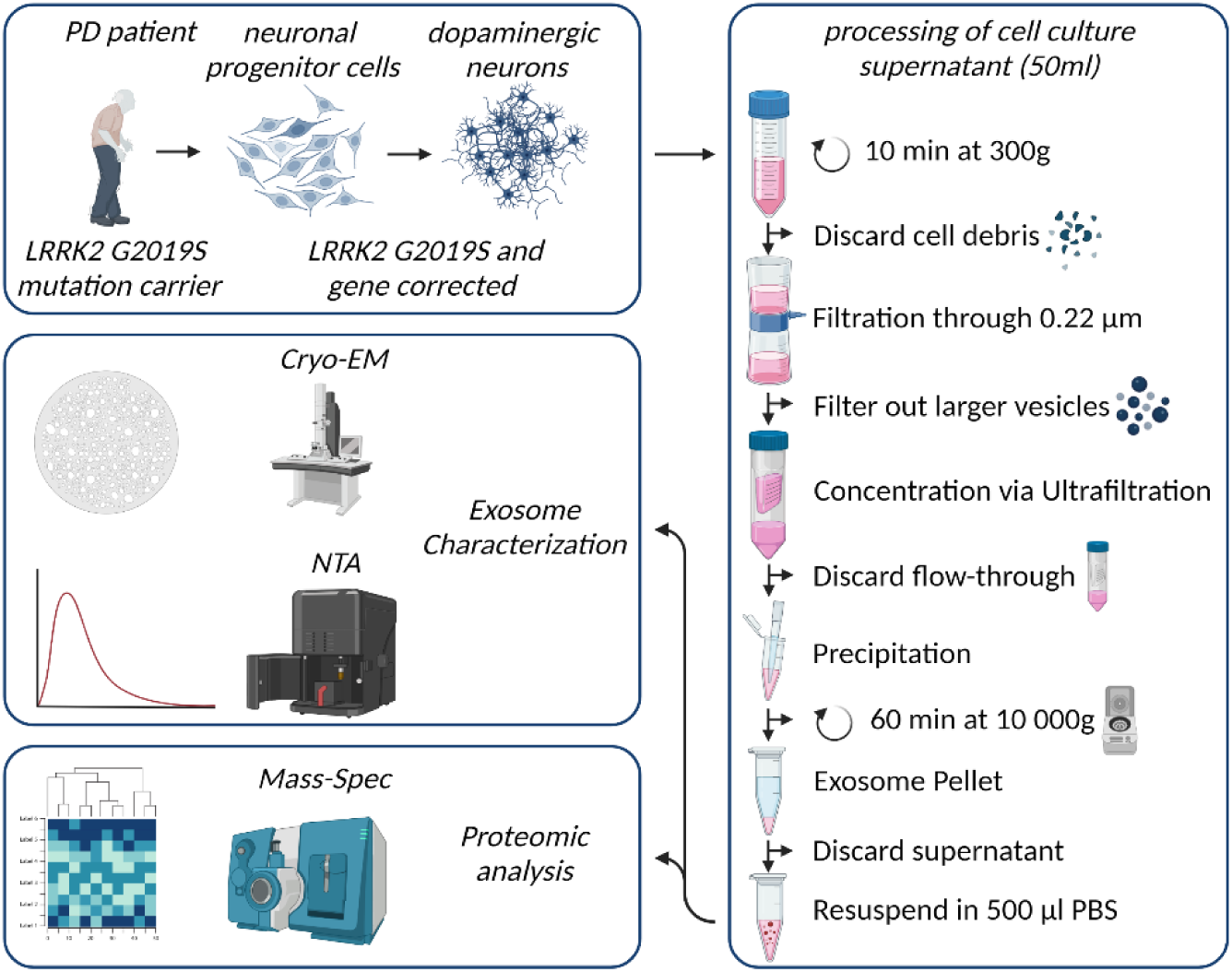
Visual representation of study design and the EV isolation protocol. Neural progenitor cells were generated previously using iPSCs derived from a female PD patient carrying the LRRK2 G2019S mutation. After induction of differentiation over a period of 14 days, cell culture supernatant was collected every three days from human dopaminergic neurons, resulting in a total of 36 ml of supernatant. Supernatant was thoroughly processed and resulting EV samples were characterized using NTA and Cryo-TEM. Finally, proteomes of EV samples and cell lysates of hDaNs were analyzed.

### Isolation of Extracellular Vesicles

To eliminate cellular debris, CCM was centrifuged at 300g for ten minutes after which it was filtered through a 0.22 µm Steriflip filter (Merck, #SCGP00525). To concentrate the CCM to 1 ml, we used the Amicon® Ultracel Centrifugal filters (Merck, #UFC901024), which were centrifuged at 2000g at 4°C for 30 minutes. After 1 ml of concentrated CCM was recovered, we transferred it to a 1.5 ml tube and added 500 µl of Total Exosome Isolation Reagent (Thermo Fischer Scientific, #4478359). After overnight incubation at 4° C, samples were centrifuged at 10 000g for one hour. Supernatant was carefully removed before tubes were centrifuged for 5 minutes at 10,000g. Finally, pellets were washed three times with 1ml PBS to remove residual traces of the isolation reagent before they were resuspended in 200ul of PBS containing cOmplete protease inhibitor (Sigma-Aldrich, #11873580001) and phosphatase inhibitor (Sigma-Aldrich, #4906837001) (Fig. 1).

### Characterization of Extracellular Vesicles

We characterized the extracellular vesicles following the minimal information for studies of extracellular vesicles guidelines (*18*): 1) particle size and concentration were analyzed using nanoparticle tracking analysis (NTA); 2) morphology of vesicles was assessed via cryo transmission electron microscopy (cryo-TEM); 3) presence of EV protein markers within the proteome of all EV samples was confirmed; and 4) gene ontology (GO) enrichment analysis for the cellular component was performed for proteins detected in all EV samples.

*Particle Size Concentration*–Measurements were conducted using the Nanosight NS300 instrument and the NanoSight NTA 3.0 0068 software provided by Malvern Panalytical in Kassel, Germany. To optimize the accuracy of the analysis, the samples were diluted in phosphate-buffered saline (PBS) at ratios ranging from 1:100 to 1:500 prior to measurement. Per measurement, five videos with a length of 60 second were recorded.

*Cryo-TEM*–Cryo Transmission Electron Microscopy was performed at the Nanoscale and Microscale Research Centre (nmRC) located in the University of Nottingham, UK. The nmRC has previously published a protocol for preparing EV samples for cryo-TEM, which was adapted for this study (*19,20*). 5 µL of EV sample was added onto each Holey carbon TEM grid for two minutes (EM resolutions, Sheffield, UK, #HC300Cu). After removal of excess solution using filters, EV samples were blotted for a second and then frozen in liquid ethane using a Gatan CP3 plunge freezing unit (Ametek, Leicester, UK). Samples were loaded to a FEI Tecnai G2 12 Bio-twin TEM. Images were obtained using an inbuilt Gatan SIS Megaview-IV digital camera at an accelerating voltage of 100 kV.

*Confirmation of presence of EV marker proteins–*We compared the relative abundancy of the EV marker proteins TSG101, CD81 and Flotillin-1 and the negative marker Calnexin in the proteome of all EV samples to their abundancy in hDaNs cell lysates. In addition, we performed a GO enrichment analysis of the most abundant proteins found in the EV samples. We therefore selected the top 100 most abundant unique protein identities in each genotype, identified the overlap between these two and analyzed the cellular components these proteomes were annotated to. GO enrichment analysis was performed using the *enrichGO* function from the ClusterProfiler package in R and is described in further details in the statistics section (*21,22*).

### DIA-based Proteomic Analysis

*Protein Extraction*–EV proteomes were quantitatively assessed by DIA-based mass spectrometry. For this purpose, the EV fractions were concentrated by lyophilization and re-dissolved in 1x Laemmli buffer. Cell lysates were collected by adding 150 µl of a simple lysing buffer (PBS containing 1% of Triton 100x, cOmplete protease and phosphatase inhibitor) to one 6-well per replicate and were further processed.

*SDS-PAGE and In-Gel Digestion*–50µg of total protein was subjected to SDS PAGE (NuPage 10% Bis-Tris gels). The electrophoresis was topped after the sample front reached one 1cm. For visualization of the protein, gels were stained by Coomassie. The lanes were excised, distained and subjected to in-gel proteolysis by a Trypsin/LysC mix (Promega) following standard protocols (*23*). Extracted and vacuum-dried peptides were subjected to an additional C18-StageTip (Thermo Fisher) pre-cleaning step (*24*). Finally, vacuum-dried samples were dissolved in 0.5% TFA and mixed with 2µL iRT standard peptide mix (Biognosis).

*LC-MS/MS Analysis*–Mass Spectrometry analysis was performed was performed on an Ultimate3000 RSLC system coupled to an Orbitrap Tribrid Fusion mass spectrometer (Thermo Fisher Scientific). Tryptic peptides were loaded onto a µPAC Trapping Column with a pillar diameter of 5 µm, inter-pillar distance of 2.5 µm, pillar length/bed depth of 18 µm, external porosity of 9%, bed channel width of 2 mm and length of 10 mm; pillars are superficially porous with a porous shell thickness of 300 nm and pore sizes in the order of 100 to 200 Å at a flow rate of 10 µl per min in 0.1% trifluoroacetic acid in HPLC-grade water. Peptides were eluted and separated on the PharmaFluidics µPAC nano- LC column: 50 cm µPAC C18 with a pillar diameter of 5 µm, inter-pillar distance of 2.5 µm, pillar length/bed depth of 18 µm, external porosity of 59%, bed channel width of 315 µm and bed length of 50 cm; pillars are superficially porous with a porous shell thickness of 300 nm and pore sizes in the order of 100 to 200 Å by a linear gradient from 2% to 30 % of buffer B (80% acetonitrile and 0.08% formic acid in HPLC-grade water) in buffer A (2% acetonitrile and 0.1% formic acid in HPLC-grade water) at a flow rate of 300 nl per min. The remaining peptides were eluted by a short gradient from 30% to 95% buffer B; the total gradient run was 120 min. Spectra were acquired in DIA (Data Independent Acquisition) mode using 50 variable-width windows over the mass range 350-1500 m/z, MS2 scan range was set from 200 to 2000 m/z.

*Data Analysis*–Data analysis was performed using DIA-NN (ver. 1.8.1) activating following options (*25*): Trypsin/P as enzyme, “FASTA digest for liberty-free search/ library generation” (database: human subset of SwissProt 2021_04, 20375 entries) and “Deep learning-based spectra, RTs an IMs predication”. In addition, the *match between runs* (MBR) option was activated. Carbamidomethylation was set as fixed modification and N-terminal methionine excision was allowed. Cross-run normalization was done RT-dependent and smart profiling was used for library generation. In addition, a heuristic model for protein inference was used. Mass accuracies and window widths were determined/ detected by the algorithm. In addition, isotopologues were considered and no shared spectra allowed. A high precision robust LC separation was assumed for quantification. The precursor FDR was set to 1%.

### Experimental Design and Statistical Rationale

From each genotype we isolated 36 ml of CCM per differentiation (3x LRRK2 G2019S, 3x LRRK2 Gene Corrected). Each of the 36 ml of CCM resulted from an individual 6-well plate of iPSC-derived human dopaminergic neurons that had been cultured as described above. All resulting EV samples were analyzed via NTA and DIA. Cryo-TEM images were made using 100 µl of one LRRK2 G2019S and one LRRK2 GC EV sample derived from an additional differentiation that was not further included in the study.

Downstream analysis on maxLFQ values (DIA-NN unique genes matrix) was performed with Perseus (ver. 1.6.7.0) (*26*). *Data were log2 transformed to facilitate the identification of proportional changes in proteins abundance between different groups. This transformation helps normalize the data and stabilize the variance, making it more suitable for subsequent statistical analysis.* After categorical annotation, only IDs with three valid values in at least one biological group were accepted. *Missing data were imputed separately for each biological group using normal distribution, the width and down shift were set to 0.8 and 1.3, respectively.* Significant changes were detected by a two-sided Student’s T-test with using a permutation-based FDR (S0=1, FDR=0.05 with 250 randomizations).

Statistical analysis of the NTA data, namely unpaired t-test on differences of particle yields and sizes between the genotypes, was performed using GraphPad Prism software, version 9.3.0. (La Jolla, CA, United States). QQ-plots were used to assess normality of the data. Mean relative abundancy of marker proteins TSG101, Flotillin-1, CD81 and Calnexin in hDaNs cell lysates was calculated. Next, for each EV sample a log2 fold change (log2 fc) value of that mean abundancy was calculated.

For further data analysis and visualization, various functions from R-Studio, version 4.3.0 were used (*22*); principal component analysis (PCA) of the six EV samples and six cell lysates was performed using the built-in *prcomp* function and using the data of proteins identified in all 12 samples. For the generation of heatmaps, proteins that were not present in all samples were excluded. The built-in *scale* function was used to normalize the data. When plotting data from hDaNs and EVs in one heatmap, intensities were normalized separately for each sample type; for downstream analysis, a fold change cut-off of >±1.5 was applied. Proteins that did not cross this threshold were not further considered, even if they passed statistical testing of differential expression. GO enrichment analysis was performed using the *enrichGO* function from the *ClusterProfiler* package (*21*); if not stated otherwise, proteins that were either up- or downregulated were analyzed separately. The p-value cut-off was set to 0.05, FDR correction was performed using the Benjamini-Hochberg procedure to correct for multiple testing and adjusted p-values will be reported (*27*). To filter out GO terms relevant to our research question, we filtered for CNS-related GO terms using the *grep* function in R and a customized list of terms (Supplemental Table S2). If applicable, the top 15 GO terms sorted by p-value were visualized using a semantic scatter plot with a maximum similarity matrix being calculated to indicate semantic proximity of identified GO terms. If applicable, we further visualized the top three GO terms and their annotated proteins in a network plot.

Finally, we used the Cytoscape software, version 3.8.2 and the String data base (*28,29*) to identify the LRRK2 interactome. The physical subnetwork was plotted and a confidence score cut-off was set at the default of 0.4.

### Semi-Automated Literature Review

To identify promising biomarker candidates, we performed a semi-automated literature research on the final set of proteins identified to be dysregulated both in EVs and cells. The search was conducted via the *entrez_search* function from the *rentrez* package in R and applied to the PubMed database to retrieve PubMed IDs associated with our identified proteins and our research question. This function returned PubMed IDs based on our defined protein list and term queries. We selected the terms “Parkinson’s disease” and “LRRK2”.

## RESULTS

### Characterization of hDaNs derived Extracellular Vesicles

NTA confirmed the presence of particles with a size of 30 to 220 nm, with peaks around 80-90 nm (Fig. 2A). Mean particle size was 98.5 nm (SD: ±1.1) in EVs from L1 GC and 103.2 nm (SD: ±8.2) in EVs from L1 G2019S. Unpaired t-test showed no statistically significant differences between the two lines (difference between means: 4.7, t = 0.99, df = 4 p = 0.376). Mean particle yield per line and batch of CCM was 3.4x10^11^ EVs (SD: 4.8x10^10^) in L1 GC and 3.6x10^11^ EVs (SD: 1.5x10^11^) in L1 G2019S. Again, unpaired t-test did not show any significant difference between lines (difference between means: 2.1x10^11^, t = 0.23, df = 4, p = 0.184) (Fig. 2B).

**FIG 2.**
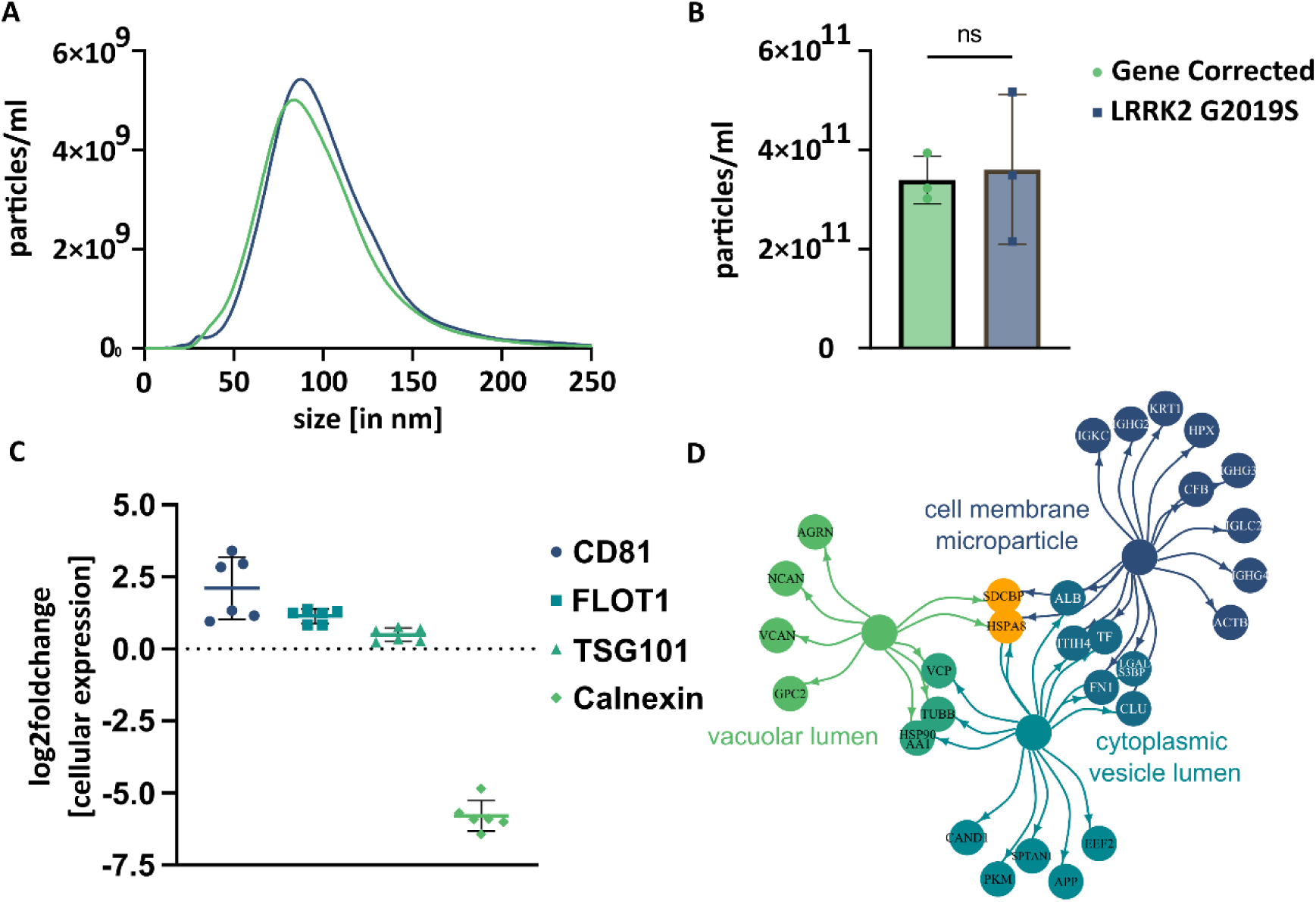
Basic characterization of EV isolated from hDaNs derived cell culture supernatant. A, NTA was performed on 3 EV samples per genotype. Size of EVs ranged from 30 to 220 nm with a peak around 80 to 90 nm. B, No significant difference of particle yield between genotypes was detected. Error bars indicate standard deviation. C, Relative abundancy of proteins in EVs is given as log fold change compared to cells. EV markers CD81, Flotillin-1 (FLOT1) and TSG101 were more abundant in EVs, while negative marker Calnexin was clearly more abundant in cells. D, GO enrichment analysis was performed for cellular component on the most abundant proteins detectable in all 6 EV samples. Depicted terms were among the top 10 significantly associated cellular components. Color of circles indicate to which term given proteins were annotated to. If a protein was annotated to two terms, depicted color is a mix of both. Orange proteins were annotated to all three terms.

We validated our isolation protocol by comparing the relative abundance of EV marker proteins, CD81, Flotillin- 1 and TSG101 and the relative scarcity of Calnexin, in the proteome of EVs and cell lysates. (Fig. 2C). Relative abundancy of CD81 (mean log2 fc compared to cells: 2.1, SD: 1.1, n = 6) and Flotillin-1 (mean log2 fc: 1.1, SD: 0.2, n = 6) was higher in EVs than in hDaNs. Abundance of TSG101was slightly higher in EVs (log2 fc: 0.5, SD: 0.2, n = 6) compared to hDaNs, while levels of Calnexin (log2 fc: -5.8, SD: 0.5, n = 6) were clearly higher in hDaNs compared to EVs.

To further assess the proteome of our EV samples and confirm their vesicular nature, we performed a GO enrichment analysis with the most abundant proteins. Of the 100 proteins selected per genotype, 77 proteins were among the most abundant in EVs from both L1 GC and L1 G2019S. Out of the 51 significantly associated GO terms (Supplemental Table S3), many were associated to either vesicles or the endosomal-lysosomal pathway. Among the top ten terms were “*cell membrane microparticle*” (associated proteins: 17/77, p<0.0001), “*cytoplasmic vesicle lumen*” (16/77, p<0.0001), and “*vacuolar lumen*” (9/77, p<0.0001), (Fig. 2D)

In Cryo-TEM, particles appeared as solitary, spherical and membrane-encapsulated structures, in line with previous reports of EVs (*30*). We did not find any morphological differences between the two genotypes (Supplemental Figure 1).

### Proteomic Analysis

*Quantitative Analysis of Proteomes*–Quantitative proteomics was performed using the data-independent analysis (DIA) approach. Three technical replicates were analyzed for EV as well as for the corresponding cell lysates of the same differentiations. Using DIA-NN, libraries were created by the extraction of pseudo-MSMS spectra directly from the DIA runs (*25*). By this approach, a total of 2634 unique proteins were consistently detectable in all 6 EV samples. The resulting heatmap separated genotypes by protein abundance (Fig. 3A). In total, 595 proteins were significantly dysregulated (n_upregulated_ = 318, n_downregulated_ = 277) in L1 G2019S compared to L1 GC (Fig. 3B and 3C). Of those, 484 passed our fold change threshold (n_upregulated_ = 275, n_downregulated_ = 209). In the cell lysates of hDaNs, a total of 5833 unique proteins were consistently detectable in all 6 samples (Fig. 3D). 3205 proteins were significantly dysregulated (n_upregulated_ = 1190, n_downregulated_ = 2015) in L1 G2019S compared to L1 GC (Fig. 3E and 3F). Of those, 1833 passed our fold change threshold (n_upregulated_ = 907, n_downregulated_ = 926).

**FIG 3.**
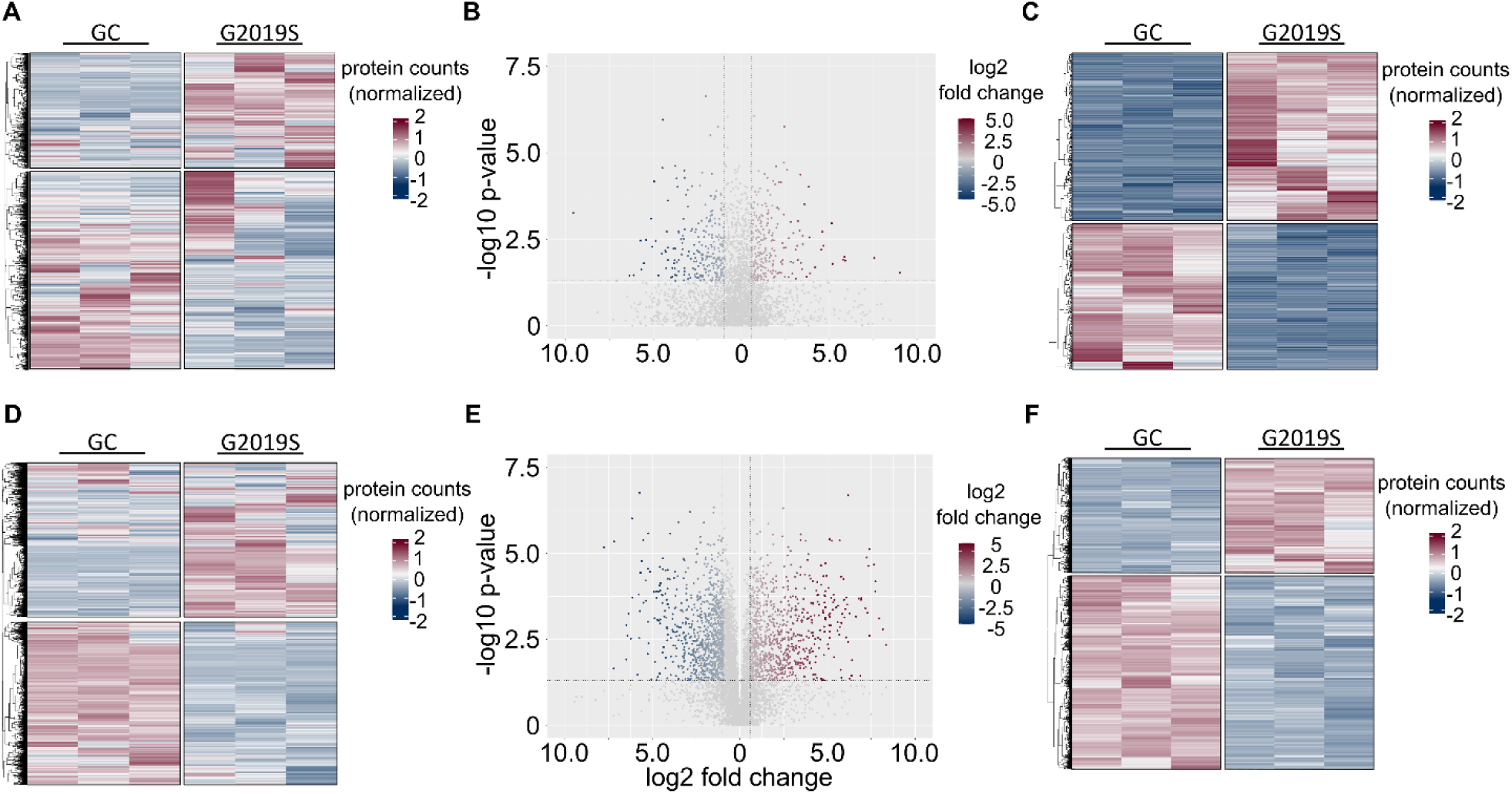
Proteomic signature of EVs and cell lysates derived from hDaNs carrying the LRRK2 G2019S mutation. A, In EVs, a total of 2634 proteins could be identified in all 6 samples. When normalizing protein counts, separation along genotype becomes apparent. B, Volcano plot visualizes log2 fold change levels of proteins found in L1 G2019S compared to L1 GC. Dotted horizontal line visualizes the p-value cut-off at 0.05, while vertical lines indicate the >±1.5 fold-change threshold. Of the 595 significantly dysregulated proteins, 484 passed this cut off. C, Heatmap of differentially expressed proteins in EV samples. D, In cell lysates of hDaNs, 5833 proteins were found. E, A total of 3205 proteins were significantly dysregulated, while 1833 passed the fold change threshold. F, Heatmap of differentially expressed proteins in cell lysates.

*Functional Enrichment Analyses*–Functional enrichment analyses of the dysregulated proteomes were performed separately for up- and downregulated proteins. For the sake of stringency, we focused on the *biological processes* significantly associated with the dysregulated proteomes. Among the significantly associated GO terms for the proteins upregulated in L1 G2019S EVs were *synapse organization*, *synapse assembly* and *neuron projection regeneration* (Fig. 4A and 4B) (Supplemental Table S4). A total of two CNS related GO terms were associated with the downregulated EV proteome, namely *regulation of postsynaptic neurotransmitter receptor activity* and *synapse organization* (Fig. 4C and 4D) (Supplemental Table S5).

**FIG 4.**
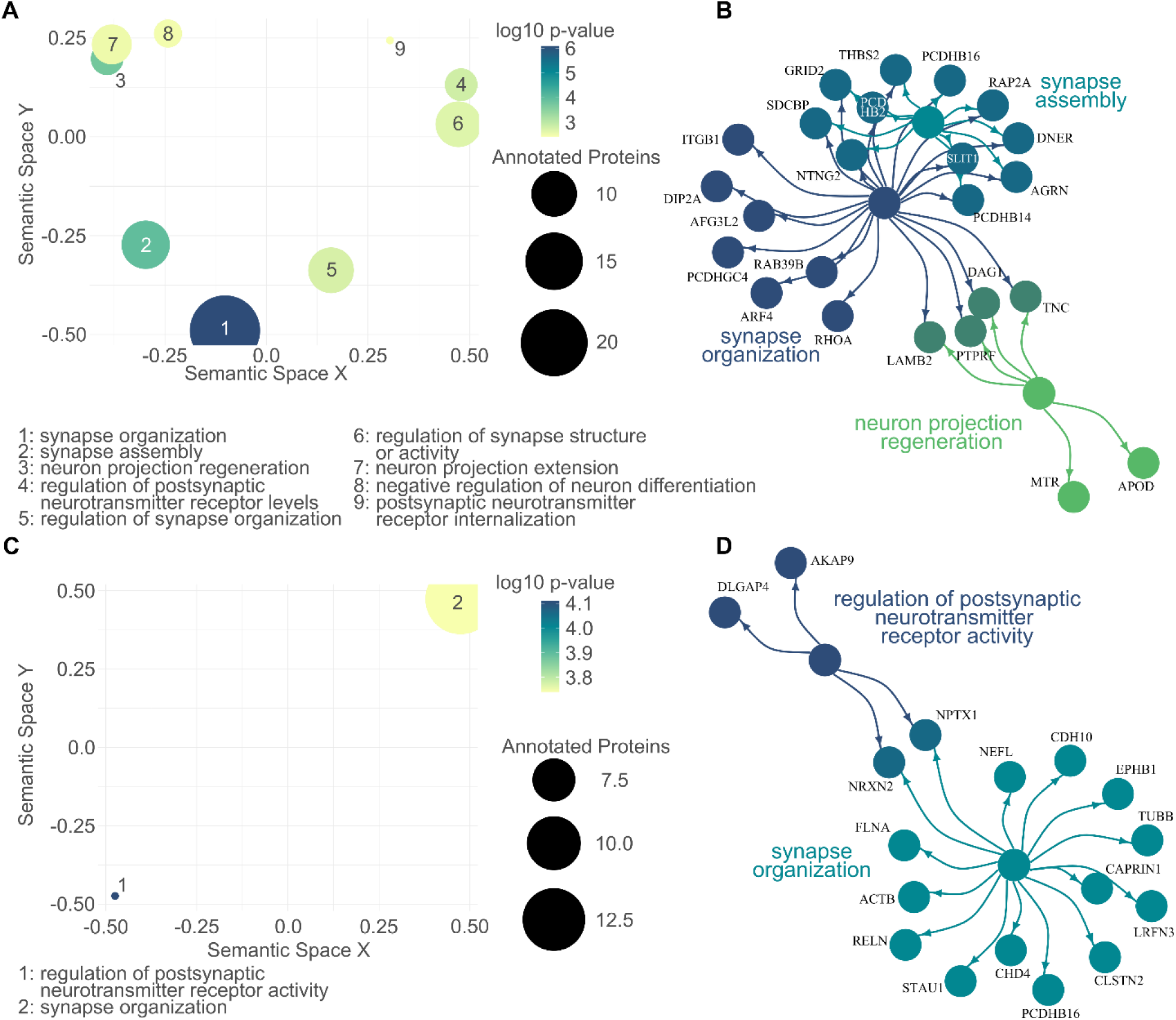
CNS related gene ontology enrichment analysis of the L1 G2019S EV proteome. A, Semantic scatter plot visualizes the identified GO terms in a two-dimensional semantic space. Terms that are semantically related are depicted more closely to each other. Size of the dots represent number of annotated proteins, while color depicts the log10 p-value. A total of nine CNS related GO terms were significantly associated with the proteins upregulated in L1 G2019S EVs. B, Of those, the top three were visualized together with their annotated proteins. Color of dots indicates the respective GO term. Proteins that were annotated to more than one term are colored in a mix of the respective colors. C, GO enrichment analysis was repeated for the proteome downregulated in L1 G2019S EVs. Here, only two CNS related GO terms were identified and plotted in D, a network plot.

As for the cellular proteome, amongst others, upregulated proteins were significantly associated to *vesicle-mediated transport in synapse*, *regulation of neurotransmitter levels* and *neurotransmitter transport* (Fig. 5A) (Supplemental Table S6). In contrast, the downregulated proteins were not significantly associated to any CNS related GO terms. We therefore did not apply the filtering step and plotted the top 15 terms resulting from the total GO enrichment analysis (Fig. 5B) (Supplemental Table S7). Among those were *RNA processing*, *RNA splicing* and *cellular nitrogen compound catabolic process*.

**FIG 5.**
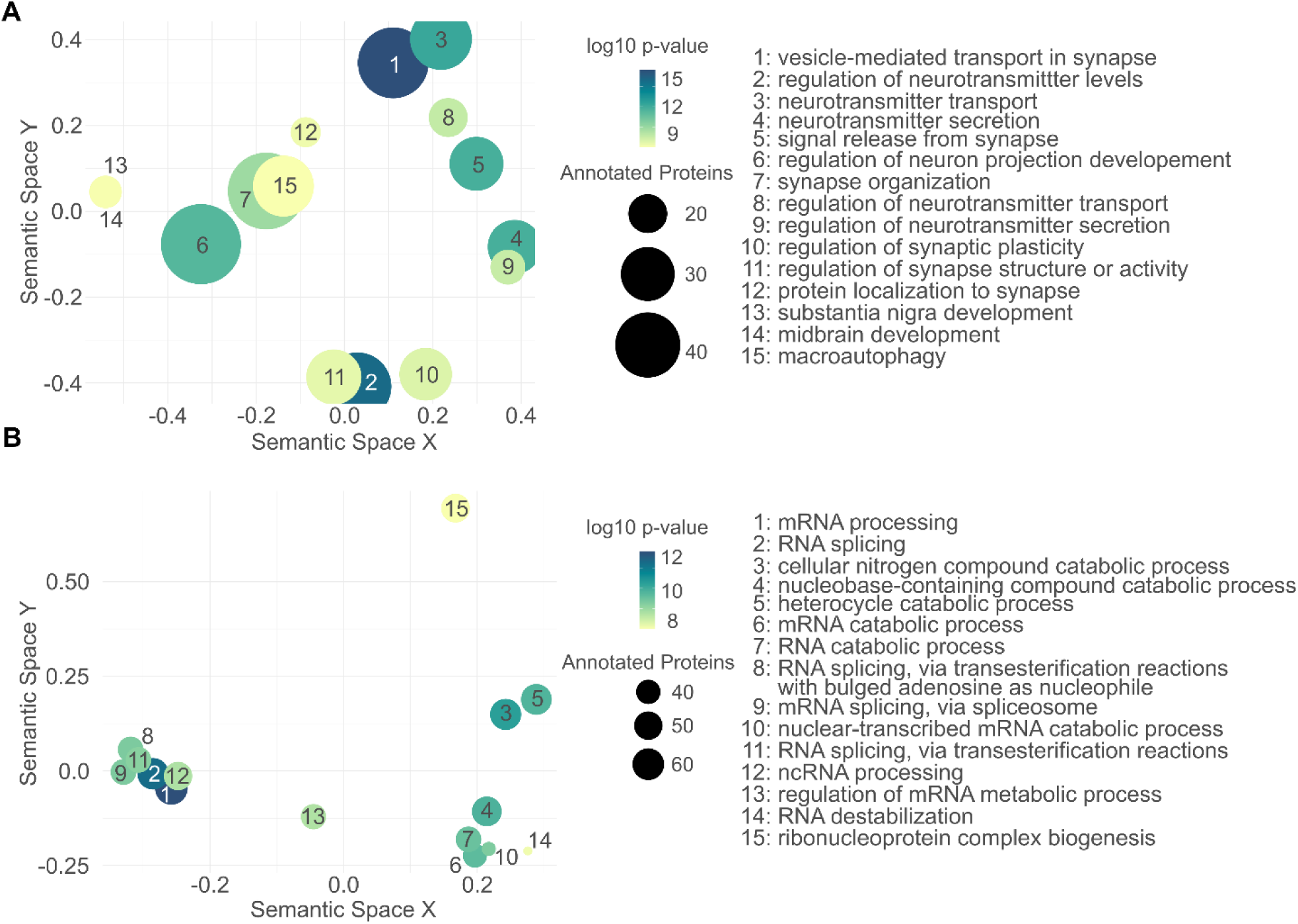
Gene ontology enrichment analysis of the L1 G2019S cellular proteome. A, Semantic scatter plot of CNS related GO terms found to be significantly associated with the upregulated proteins in L1 G2019S hDaNs. B, In contrast to the upregulated proteins, no CNS related GO term was significantly annotated to the downregulated proteome. Instead, the unfiltered GO terms are plotted, most of which seem to indicate changes in RNA related processes.

*Analysis of commonly dysregulated proteins*–As a final step, we analyzed the proteins that were dysregulated both in the proteome of EVs and hDaNs cell lysates to increase the robustness of our findings. First, we performed PCA using data from all proteins identified in both EVs and cell lysates (Fig. 6A). PC1 explained 53.1% of the variability and separated EVs from cell lysates, while PC2 explained 13.5% of the variability and separated genotypes. Notably, data from the two cellular proteomes, i.e., L1 GC and L1 G2019S, clustered closer together compared to the EV proteomes. Next, we looked at the overlap of dysregulated proteins (Fig. 6B). Of the 484 dysregulated proteins in EVs and 1833 dysregulated proteins in hDaNs, a total of 123 proteins were found to be dysregulated in both. 34 proteins were found to be upregulated, while 28 were downregulated in both sample types. The remaining 61 proteins were were dysregulated in opposite directions, e.g. upregulated in cells but downregulated in EVs and vice versa (Fig. 6B and &C). Using Cytoscape and the String data base we identified and visualized those proteins among the 123 that were described to be within the LRRK2 interactome (Fig. 6D). Finally, we performed a GO enrichment analysis using all 123 proteins as an input and identified, amongst others, the GO terms *synapse organization*, *cell migration in hindbrain* and *regulation of synapse organization* to be significantly annotated (Fig. 6E and F) (Supplemental Table S8).

**FIG 6.**
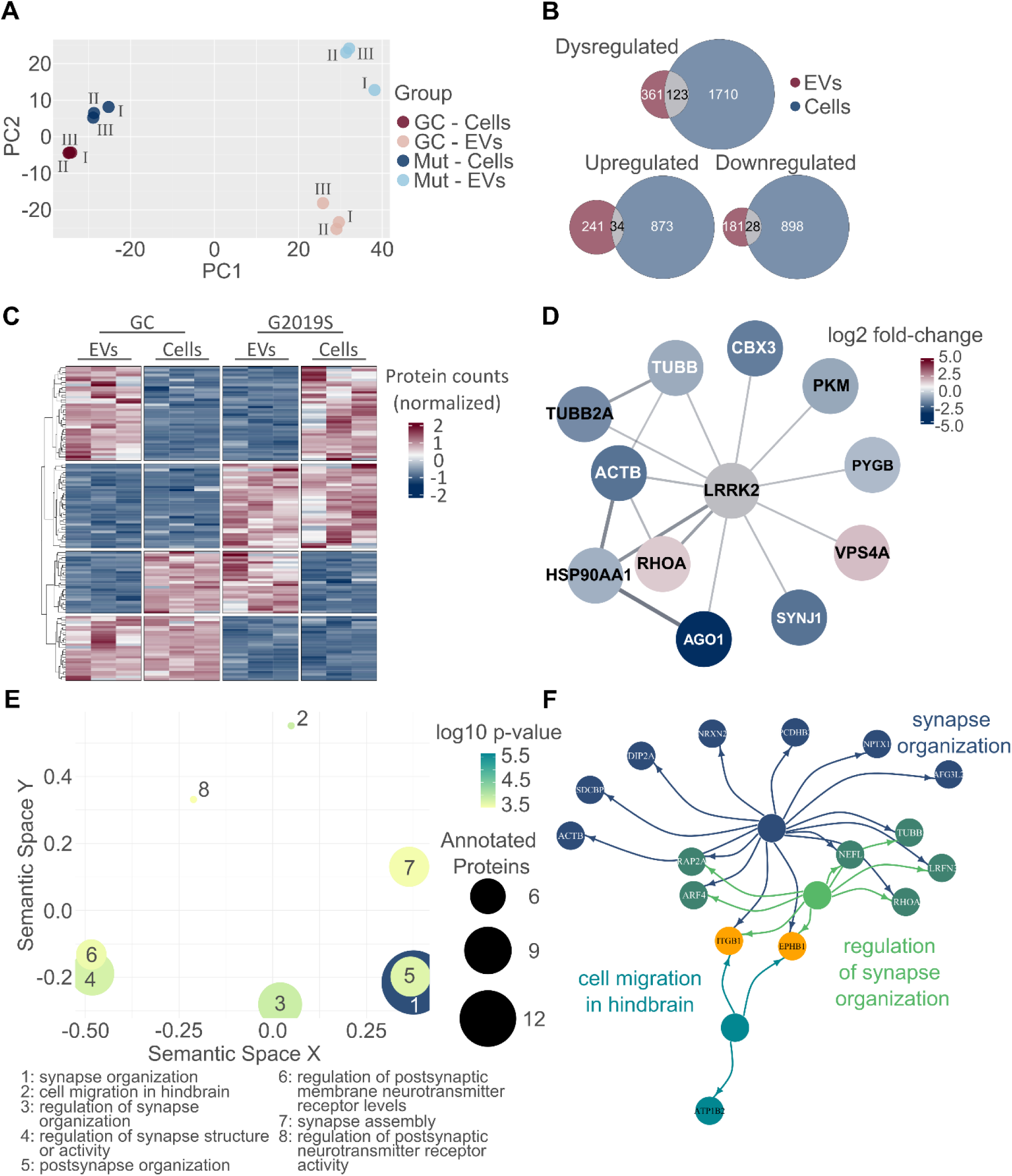
Analysis of commonly dysregulated proteins. A, PCA analysis was performed on all proteins identified in both EVs and cell lysates. EVs are clearly separated from cell lysates along PC1, while PC2 separates along genotype. Notably, difference between proteomes seems to be larger in EVs compared to cells. B, A total of 123 proteins were dysregulated on both EVs and cells as indicated by the overlap colored in grey. Direction of dysregulation was the same in EVs and cells for 34 and 28 proteins, respectively. C, The heatmap visualizes the normalized protein intensities in each technical replicate and in EVs and cell lysates. Normalization was performed for each compartment (EVs vs cell lysates) separated. Interestingly, a considerable fraction of proteins was not dysregulated in the same direction when comparing the cellular and extracellular proteome of the same genotype. D, Using Cytoscape and the String database, a total of 11 proteins appears to be within the LRRK2 interactome. Colors are based on the fold-change expression in G2019S EVs E, GO analysis was performed on all 123 proteins dysregulated in both EVs and hDaNs. F, The top three hits are visualized together with their annotated proteins. Again, the color of circles indicates towards which of the GO terms a given protein was annotated. Proteins annotated to two terms are colored in a mixture of the term’s color while orange indicates annotation towards all three terms.

### Semi-automated literature review

Out of the 123 proteins, for 76 proteins we found at least one PubMed ID when using the protein name in combination with “*AND Parkinson’s disease*”, totaling up to 608 publications. For the combination of a protein and the term “*AND LRRK2*” we found 57 publications for 22 proteins, all of which were also identified using the “*AND Parkinson’s disease*” term (Supplemental Fig. 2). Out of these 22 proteins, we identified 10 interesting candidates based on the available literature (Table 1).

**TABLE 1.** Overview of functionally relevant biomarker candidates.

| Protein Name | Gene Name | Log2 fc EVs | Log2 fc hDaNs | String LRRK2 Interactome | Function (38,39) | Context |
| --- | --- | --- | --- | --- | --- | --- |
| Actin Beta | ACTB | -2.3 | 6.6 | yes | crucial and ubiquitously expressed protein of the cytoskeleton | enriched in synapses, role of LRRK2 in cytoskeleton reorganization (44) |
| Argonaute RISC Component 1 | AGO1 | -3.6 | -1.6 | yes | RNA interference, e.g. repression of mRNA translation, binding to miRNAs | dysregulated RNA processing in PD, LRRK2 driven miRNA signatures (46,47) |
| Annexin A1 | ANXA1 | 2.4 | 2.1 | no | inhibition of phospholipase A2 | role in neuroinflammation (51) |
| CD44 | CD44 | 3.8 | 3.2 | no | cell-surface protein, role in cell-cell interaction and migration | role in neuroinflammation, upregulated in CSF of LRRK2 PD (53, 64) |
| Galactosylceramidase | GALC | -3.9 | -1.0 | no | lysosomal protein involved in sphingolipid metabolism | related to GBA, influence on lysosomal membranes (56) |
| Peroxiredoxin 2 | PRDX2 | 0.9 | -1.0 | no | antioxidant enzyme | Phosphorylated by LRRK2, leading to loss of dopaminergic neurons (62) |
| Ras Homolog Family Member A | RHOA | 1.0 | 1.0 | yes | small GTPases, promotes cytoskeleton reorganization | Remodeling of synaptic cytoskeleton, linked to damage of dopaminergic neurons (40,41) |
| Sonic Hedgehog Signaling Molecule | SHH | 1.0 | 2.9 | no | key signaling molecule | Disruption of SHH signaling in LRRK2 as a mechanism of neuronal vulnerability (49,50) |
| Synaptojanin 1 | SYNJ1 | -2.3 | 1.3 | yes | synaptic transmission and membrane trafficking | Phosphorylated by LRRK2, deregulating synaptic vesicle trafficking (35) |
| Tubulin Beta Class I | TUBB | -0.9 | 0.4 | yes | structural component of microtubules | LRRK2 leading to microtubule instability (36) |

## DISCUSSION

In the present study, we aimed to identify protein-based biomarker candidates, that are functionally relevant in the context of PD and are linked to LRRK2. This goal represents an important step towards developing targeted therapies, especially with LRRK2 inhibitors rapidly approaching clinical trials (*10*). We isolated EVs from hDaNs carrying the LRRK2 G2019S mutation and an isogenic, gene-corrected control and used DIA-proteomics to analyze the genotype-specific cellular and extracellular proteomic signatures. Finally, we performed a gene ontology analysis for both the cellular and extracellular proteomes and thoroughly reviewed the literature to identify promising candidates for future pre-clinical or clinical biomarker studies.

As a result of our GO enrichment analysis on dysregulated proteins found both in EVs and cell lysates, we identified two major biological processes to be dysregulated in the LRRK2 G2019S cell model, one of which was the structural and functional integrity of synapses. Synapses have extensively been described to potentially play a critical role in the development of neurodegenerative diseases, including LRRK2-linked PD. For example, one study showed LRRK2-dependent regulation of the pre- and postsynaptic morphology via the interaction with microtubule-associated-protein-like targets (*31*). LRRK2 was also shown to interfere with endocytosis of synaptic vesicles, thus affecting neurotransmission (*32*) and elevated LRRK2 kinase activity was shown to affect striatal dopamine release and uptake in a LRRK2 G2019S mouse model (*33*). Along these lines of evidence is the identification of dysregulated levels of Synaptojanin 1 (SYNJ1), Tubulin Beta Class I (TUBB), cytoskeletal protein Actin ß (ACTB) and Ras Homolog Family Member A (RHOA) in our cell model.

SYNJ1, which is involved in synaptic transmission and membrane trafficking, interacts with several proteins playing a key role in synaptic vesicle recycling processes (*34*). Excitingly, it was described to be directly phosphorylated by LRRK2, suggesting a close interplay of these two proteins in deregulating trafficking of synaptic vesicles (*35*). TUBB, which forms a structural component of microtubules, was also shown to interact with LRRK2, resulting in affection of microtubule stability (*36*). As neuronal cells are highly dependent on an effective intracellular transport provided by microtubules, their disruption is likely an early event in many neurodegenerative disorders (*37*). RHOA on the other hand, likely controls the actin cytoskeleton’s rearrangement and is crucial for coordinating the formation and remodeling of synapses(*38–40*). More importantly, multiple studies have drawn a connection between increased RHOA activity and PD-linked phenomena, such as damage of dopaminergic neurons or upregulation of alpha-synuclein (*41*). In addition, an interaction with LRRK2 was discussed, although the results remained inconclusive (*42*). Finally, ACTB represents a ubiquitously expressed protein and is a crucial component of the cytoskeleton(*38,39*). As such, it is typically enriched in synapses, where it influences synaptic shape, size and neurotransmitter release (*43*). Interestingly, in a G2019S drosophila model and using RNA sequencing, ACTB was identified among the most significant dysregulated gene nodes, suggesting a critical role of LRRK2 in actin cytoskeleton reorganization (*44*).

The other biological process we identified through GO enrichment analysis was RNA processing. The disturbance of RNA processing is being discussed as both a potential cause of neurodegenerative diseases and a target for therapeutic interventions (*45*). RNA binding proteins were shown to interfere with several steps of RNA metabolism, including splicing, mRNA transport, translation and degradation. Changes such as dysregulated expression, cellular mislocalization and aggregation of RNA binding proteins were suggested to result in impaired RNA metabolism in neurodegenerative diseases, although the precise mechanisms of how this leads to neurodegeneration are not fully understood (*45*). Further, we recently showed that LRRK2 mutation carriers display distinct miRNA patterns, connecting LRRK2-driven PD to a disturbed RNA metabolism (*46*). In this context, the downregulation of AGO1 observed in our study in both hDaNs and EVs becomes of particular interest. AGO1, which, together with other proteins, forms the RNA-induced silencing complex, binds to microRNAs, and then regulates the translation of mRNAs, thereby playing a crucial role in transcriptional silencing (*38,39*). In a drosophila model it was shown that LRRK2 not only associates with AGO1, but that LRRK2 G2019S negatively regulates AGO1 expression levels (*47*). In our cell model, this downregulation seems to be measurable not only in hDaNs but also EVs, therefore confirming previous work on RNA processing and LRRK2.

Next to the above-mentioned proteins SYNJ1, TUBB, RHOA, ACTB and AGO1, we identified another five proteins from our data set that were dysregulated both in the cellular and extra-cellular proteome and appeared as promising biomarker candidates based on probable functional relevance and previous appearances in publications related to PD or LRRK2. Particularly exciting to us was the identification of upregulated levels of sonic hedgehog signaling molecule (SHH) in both EVs and hDaNs. SHH was first described to play an important role in sustaining the chemical and cellular integrity of nigrostriatal circuits over ten years ago (*48*). Since then, using cellular models of both sporadic and familial PD, including LRRK2, we have learned that there is a close interplay of increased SHH activity and ciliary dysfunction(*49*). Ciliary defects were also shown in LRRK2 mutant mice, including the G2019S variant. Disruption of the SHH signaling pathway, which requires intact cilia, might be a mechanism of neuronal vulnerability in PD patients with LRRK2 mutations and monitoring extracellular levels of SHH via quantification from EVs might be an exciting approach to e.g. monitor treatment efficacies (*50*).

Annexin A1 (ANXA1) is a membrane-bound protein and has anti-inflammatory properties, likely via the inhibition of phospholipase A2 (*38,39*). Missense variants of ANXA1 were discussed to cause genetic PD and it was hypothesized that ANXA plays a role in impaired clearance of accumulated alpha-synuclein via microglial defects and neuroinflammation (*51*). Interestingly, growing evidence suggests a link between LRRK2 and inflammation, both in the periphery and the central nervous system. LRRK2 variants were shown to modulate the risk for Crohn’s disease, a chronic inflammatory bowel disease and LRRK2 kinase activity was shown to influence microglial activation and pro-inflammatory cytokine production (*52,53*). In the same context, the upregulation of CD44 seems intriguing, as this protein is heavily involved in cell-cell interactions, especially in the context of inflammation (*38,39*). Downregulation of CD44 was shown to decrease microglia-driven neuroinflammation and reduce loss of dopaminergic neurons in CD44 knockout mice (*54*). Excitingly, a large proteomics study on CSF from both sPD and fPD patients carrying a LRRK2 mutation already revealed upregulated levels of CD44 expression compared to those in healthy controls, therefore being in line with our findings (*55*).

Galactosylceramidase (GALC) was another intriguing candidate we identified within our data set. This lysosomal protein represents an important component in the glycosphingolipid metabolism and is therefore biochemically closely related to the glucocerebrosidase (GBA) [(*38,39,56*). GBA, in turn, is particularly known for its significance in the context of PD, as a multitude of genetic variants modulate the risk of developing the disease (*57*). A recent study proposed GALC to also affect the risk of PD and suggested that its dysregulated activity might alter the composition of lysosomal membranes (*56*), which due to its involvement in the endolysosomal system links GALC directly to LRRK2 (*58*). Additionally, in a small case report, patients with dual mutations in both GBA and LRRK2 displayed increased GALC activity (*59*). Based on our data we propose that total protein levels of GALC could also be altered in patient-derived materials, including brain-derived EVs.

Finally, we found dysregulated levels of Peroxiredoxin 2 (PRDX2), which is an antioxidant enzyme that contributes to the cellular protection against damage from free radical oxygen species (ROS) (*38,39*). It is well established that regulation of ROS, which physiologically occur during the mitochondrial electron transfer chain, is essential to cellular homeostasis. Further, several PD-related genes encode proteins tightly involved in regulation of mitochondrial integrity and oxidative stress, including LRRK2 (*60*). In this context, PRDX2 overexpression was shown to be protective in a cell model of PD (*61*). Excitingly, it was also shown to be phosphorylated by LRRK2, leading to the loss of dopaminergic neurons (*62*).

In summary, in the present study we used iPSC-derived hDaNs and their EVs to characterize the proteomic changes in the context of the LRRK2 G2019S mutation. While previous studies on the proteome of urinary EVs and the CSF of LRRK2 PD patients have been performed, to the best of our knowledge this was the first study that directly characterized the proteome of dopaminergic neurons carrying a LRRK2 mutation and their EVs (*55,63,64*). We identified a plethora of dysregulated proteins and gene ontology enrichment analysis highlighted pathways that are functionally relevant for neurodegeneration. Finally, we focused on proteins that showed dysregulation on both a cellular and extra-cellular level, intending to increase robustness of our findings. We thoroughly discussed ten promising protein biomarker candidates, their functional relevance in neurodegeneration and PD as well as their connection to LRRK2, thereby providing researchers with a solid foundation for future biomarker studies.

## DATA AVAILABILITY

The generated and analyzed data are available from the corresponding author upon reasonable request.

---

**Supplemental FIG 1.**
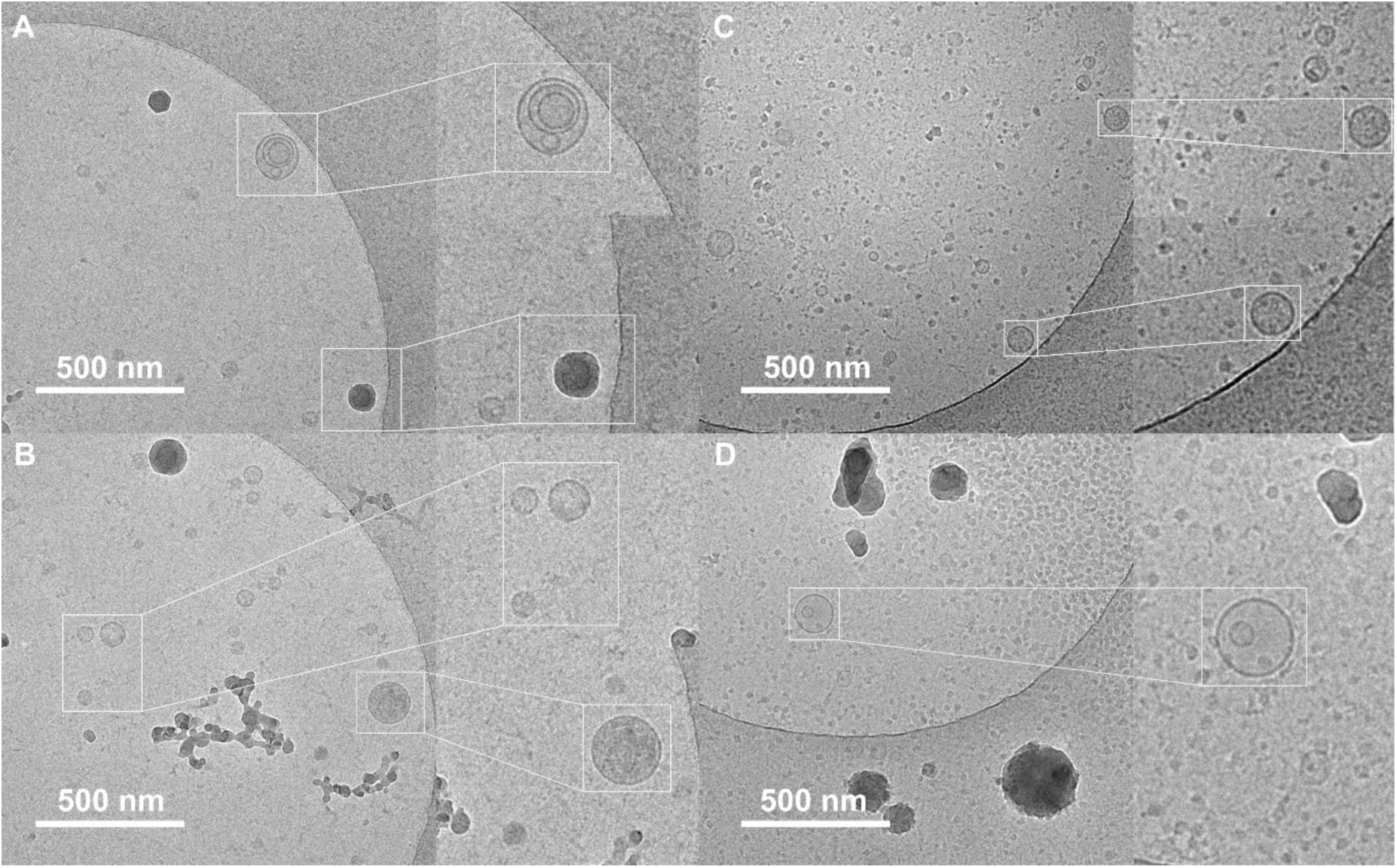
Cryo-TEM images of vesicles isolated from hDaNs. A through B, Images were taken from vesicles isolated from L1 GC as well as from C through D, L1 G2019S. Vesicles appeared as solitary, spherical and membrane-encapsulated structures. No morphological difference between genotypes were observed.

**Supplemental FIG 2.**
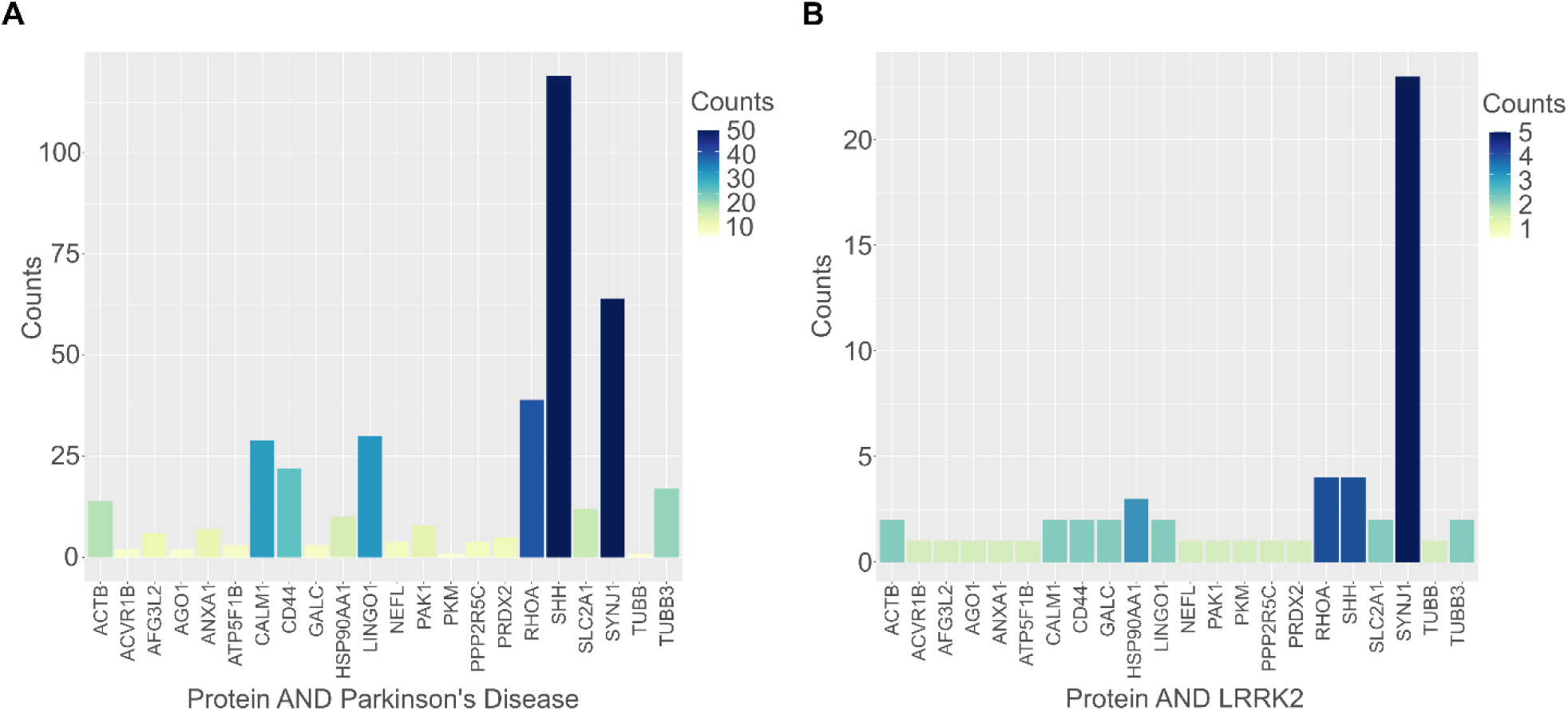
Literature review for dysregulated proteins. A, For 22 of the 123 proteins dysregulated in both EVs and hDaNs, there was at least one publication connecting it to both Parkinson’s disease and B, LRRK2.

